# Displacement-Optimized Tanglegrams for Trees and Networks

**DOI:** 10.1101/2025.11.26.690634

**Authors:** Daniel H. Huson

## Abstract

Phylogenetic trees and networks play a central role in biology, bioinformatics, and mathematical biology, and producing clear, informative visualizations of them is an important task. Tanglegrams, which display two phylogenies side by side with lines connecting shared taxa, are widely used for comparing evolutionary histories, host-parasite associations, and horizontal gene transfer. Existing layout algorithms have largely focused on trees and on minimizing the number of inter-taxon edge crossings.

We introduce displacement-optimized tanglegrams (DO-tanglegrams), a new approach that applies equally to trees and rooted phylogenetic networks. Our method explicitly minimizes taxon displacement - the vertical misalignment of corresponding taxa across the two sides - and reticulate displacement - the vertical distance spanned by reticulation edges within a network. We formalize one-sided and two-sided optimization problems, show that exact minimization is computationally intractable, and propose a heuristic that combines exhaustive local search with simulated annealing. The algorithm naturally accommodates unresolved nodes (multifurcations or multicombinations) and missing taxa. We have implemented the DO-tanglegram algorithm in SplitsTree. We compare our implementation against the cophylo R-package on a collection of synthetic trees, and against the NN-tanglegram algorithm in Dendroscope on a collection of synthetic networks. The results indicate that DO-tanglegram performs significantly better than cophylo on trees and than NN-tanglegram on networks.

**Graphical abstract:** 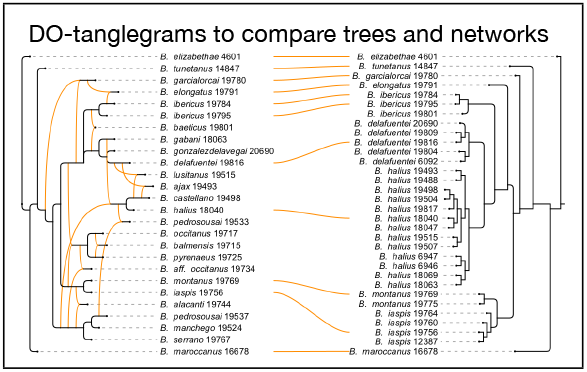

## 1 Introduction

Tanglegrams, which display two rooted trees or networks side by side with lines connecting shared (or matched) taxa, are a widely used tool in phylogenetics and comparative biology. They were popularized in the context of reconciling host–parasite associations and comparing evolutionary histories [Page, 1994, Charleston, 1998], where minimizing the number of inter-taxon edge crossings is a natural criterion for obtaining clear and interpretable drawings.

Formally, a tanglegram (for phylogenetic trees) consists of two rooted trees defined on overlapping sets of taxa, drawn facing each other, with lines or curves connecting leaves represented matching taxa, called inter-taxon edges. The internal structure of each tree is drawn planarly, so any crossings in the figure occur only among the inter-taxon edges. The tanglegram crossing minimization problem asks for leaf orders on the two sides (respecting the ancestor–descendant relationships of the trees) that minimize the number of inter-taxon edge crossings. In the one-sided version, the leaf order of one tree is fixed and only the other tree may be permuted, while in the two-sided version, both trees may be reordered subject to their constraints.

In addition to minimizing crossings, an alternative objective for tree tanglegrams is to minimize the *taxon displacement* between the two leaf orders. This measure is defined as the sum of the absolute differences between the vertical positions of corresponding leaves, and in the drawing corresponds to minimizing the total vertical displacement of the inter-taxon edges. Whereas crossing minimization emphasizes reducing visual clutter, taxon displacement emphasizes vertical alignment of matching taxa, and can yield layouts that better highlight similarities in the hierarchical structure of the two trees. This criterion was considered in Venkatachalam et al. [2010], where it is referred to as the Spearman footrule distance.

For trees, the one-sided crossing minimization problem can be solved using a polynomial *O*(*n* log *n*) algorithm [Fernau et al., 2010, Venkatachalam et al., 2010]. The more general two-sided version, where both trees can be permuted, is NP-complete. Bansal et al. [2009] introduced generalized binary tanglegrams and provided algorithms and applications motivated by comparative genomics.

Scornavacca et al. [2011] extended the concept of tanglegrams from pairs of rooted phylogenetic trees to pairs that involve rooted phylogenetic networks. Their NN-tanglegram method (implemented in Dendroscope) first computes a distance matrix *H* over the full taxon set that captures the topology of the two networks, and then applies the Neighbor-Net algorithm to *H* to derive a leaf ordering used to draw the networks. They prove that this approach is guaranteed to produce a layout with zero inter-taxon edge crossings whenever such an embedding exists. In practice, their implementation incorporates an additional heuristic step, not described in detail, that attempts to further reduce the number of inter-taxon edge crossings, without regard for the layout of the reticulate edges. NN-tanglegrams are not restricted to backbone-based layouts (as introduced below).

When moving from trees to rooted phylogenetic networks, the presence of reticulation edges introduces additional layout challenges. In this context, an analogue of taxon displacement is to minimize the *reticulate displacement*, defined as the sum of the vertical distances between the endpoints of reticulate edges in a left–to–right drawing. Whereas taxon displacement measures how well corresponding taxa align across the two sides of a tanglegram, reticulate displacement measures how nearly horizontal the reticulation edges are within a single network. Minimizing this displacement improves the readability of network tanglegrams by reducing the visual distortion caused by steeply slanted reticulation edges, and thus provides a complementary optimization criterion alongside inter-taxon edge crossing minimization.

Let *N*_1_ and *N*_2_ be two rooted phylogenetic networks (or trees) on taxon sets *X*_1_ and *X*_2_. For the purposes of this paper, a *tanglegram* consists of a left-to-right drawing of the first network *N*_1_ and a right-to-left drawing of the second network *N*_2_, separated by a rectangular region that contains lines or curves connecting any two leaves *ℓ*_1_ in *N*_1_ and *ℓ*_2_ in *N*_2_ that are labeled by matching taxa, as illustrated in Fig. 1.

**Figure 1.**
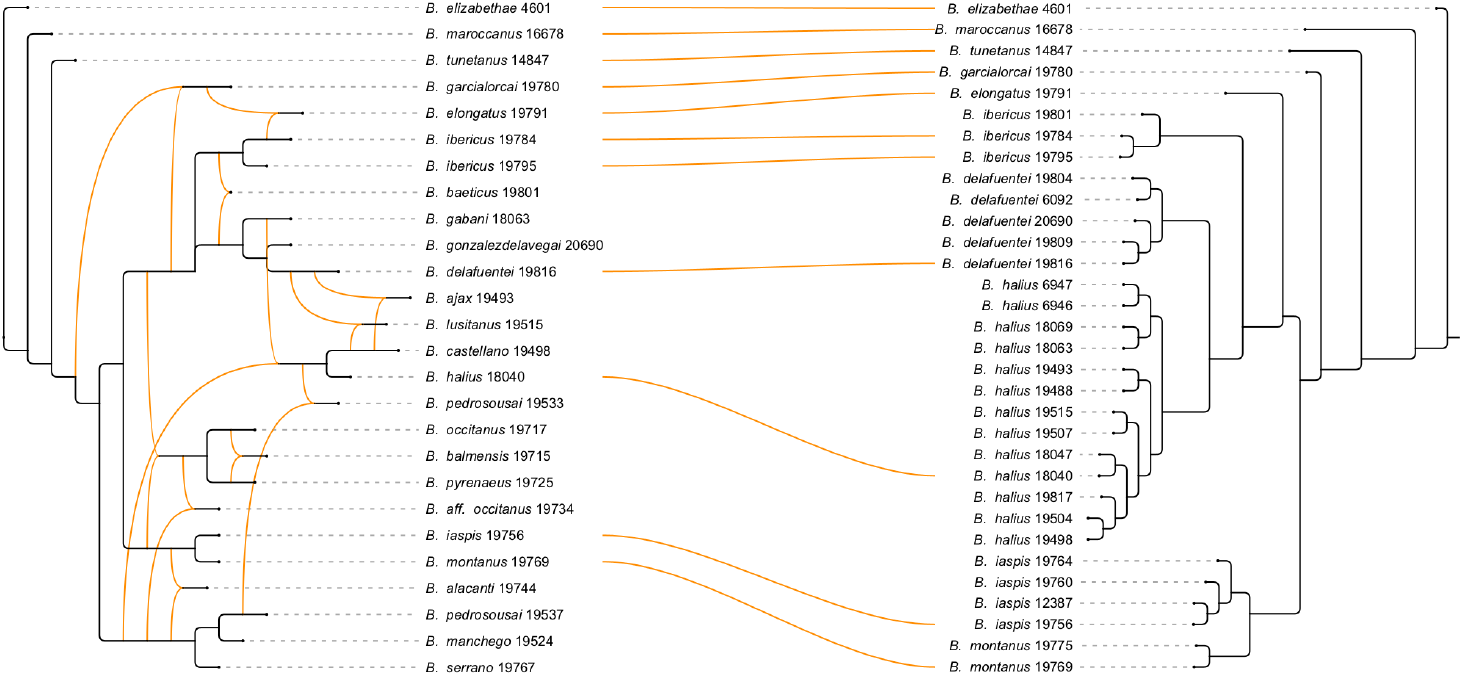
Example of a tanglegram. On the left we show a published rooted phylogenetic network on scorpions [Blasco-Aróstegui et al., 2025a] computed from gene trees using PhyloFusion [Zhang et al., 2025], and on the right, we show a phylogenetic tree from another publication [Blasco-Aróstegui et al., 2025b].

In this work we address the problem of computing a tanglegram for any two such networks, with the goal of obtaining a visually effective layout. The key idea is to explicitly minimize both the *taxon displacement* and the *reticulate displacement* of the two networks. We consider both the one-sided and two-sided variants of the tanglegram optimization problem, and we call the resulting visualization a *displacement-optimized tanglegram* (DO-tanglegram). The one-sided optimization problem is computationally hard in the case of a network [Huson, 2025], and the two-sided problem is computationally hard even in the special case where both inputs are binary trees [Fernau et al., 2010]. These hardness results motivate the development of efficient heuristics; accordingly, we propose a practical DO-tanglegram algorithm and evaluate its performance using synthetic data.

## 2 Results

Several methods have been developed for drawing tanglegrams, ranging from early tools for host–parasite cophylogenies to recent heuristics for phylogenetic networks. These approaches differ in the optimization objectives they pursue (minimizing inter-taxon edge crossings, minimizing vertical displacement, or heuristically balancing both), in the type of input they accept (binary vs. non-binary trees, identical vs. differing taxon sets, or general networks), and in whether they provide an integrated visualization environment. Table 1 summarizes the capabilities of the most relevant methods, highlighting the variants of the problem addressed and which criteria are explicitly optimized.

**Table 1:**
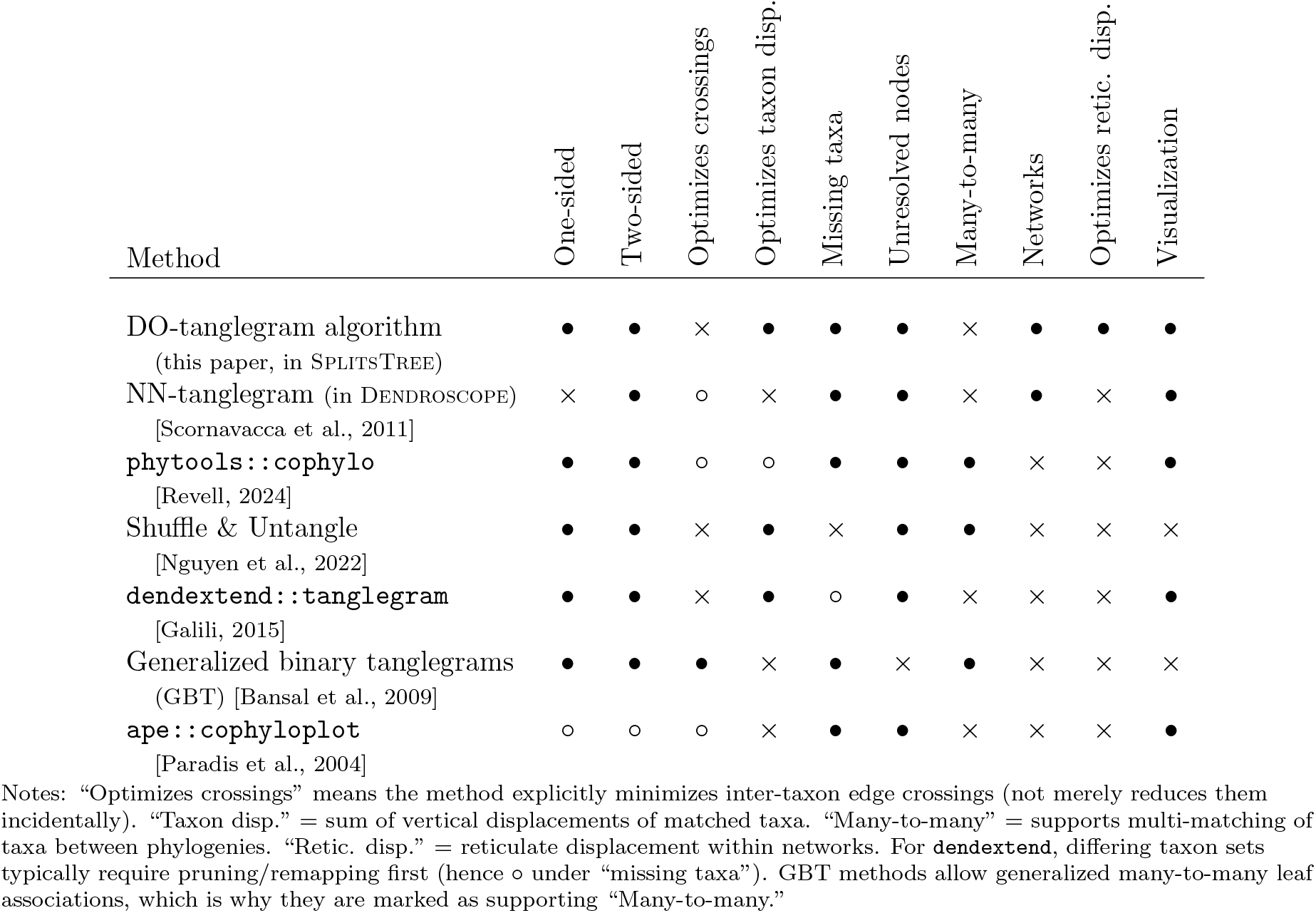
Feature comparison of tanglegram methods. For popular methods (rows), we indicate the presence of features (columns) as follows: • = supported, ∘ = partial/indirect, × = not supported.

| Method | One-sided | Two-sided | Optimizes crossings | Optimizes taxon disp. | Missing taxa | Unresolved nodes | Many-to-many | Networks | Optimizes retic. disp. | Visualization |
| --- | --- | --- | --- | --- | --- | --- | --- | --- | --- | --- |
| DO-tanglegram algorithm<br>(this paper, in SPLITSTREE) | $\bullet$ | $\bullet$ | × | $\bullet$ | $\bullet$ | $\bullet$ | × | $\bullet$ | $\bullet$ | $\bullet$ |
| NN-tanglegram (in DENDROSCOPE)<br>[Scornavacca et al., 2011] | × | $\bullet$ | $\circ$ | × | $\bullet$ | $\bullet$ | × | $\bullet$ | × | $\bullet$ |
| phytools::cophylo<br>[Revell, 2024] | $\bullet$ | $\bullet$ | $\circ$ | $\circ$ | $\bullet$ | $\bullet$ | $\bullet$ | × | × | $\bullet$ |
| Shuffle & Untangle<br>[Nguyen et al., 2022] | $\bullet$ | $\bullet$ | × | $\bullet$ | × | $\bullet$ | $\bullet$ | × | × | × |
| dendextend::tanglegram<br>[Galili, 2015] | $\bullet$ | $\bullet$ | × | $\bullet$ | $\circ$ | $\bullet$ | × | × | × | $\bullet$ |
| Generalized binary tanglegrams<br>(GBT) [Bansal et al., 2009] | $\bullet$ | $\bullet$ | $\bullet$ | × | $\bullet$ | × | $\bullet$ | × | × | × |
| ape::cophyloplot<br>[Paradis et al., 2004] | $\circ$ | $\circ$ | $\circ$ | × | $\bullet$ | $\bullet$ | × | × | × | $\bullet$ |

Notes: “Optimizes crossings” means the method explicitly minimizes inter-taxon edge crossings (not merely reduces them incidentally). “Taxon disp.” = sum of vertical displacements of matched taxa. “Many-to-many” = supports multi-matching of taxa between phylogenies. “Retic. disp.” = reticulate displacement within networks. For **dendextend**, differing taxon sets typically require pruning/remapping first (hence $\circ$ under “missing taxa”). GBT methods allow generalized many-to-many leaf associations, which is why they are marked as supporting “Many-to-many.”

We have implemented our DO-tanglegram heuristic in a new release of SplitsTree [Huson and Bryant, 2024], where it can be run in either one-sided or two-sided mode, optimizing taxon displacement and/or reticulate displacement in one or both phylogenies.

### DO-tanglegrams on trees

To demonstrate performance on trees, we compared our DO-tanglegram implementation in SplitsTree with a recently updated and widely used method for trees, the cophylo function in the R package `phytools` [Revell, 2024].

The documentation of cophylo includes an example dataset of fig wasps and their parasites [López-Vaamonde et al., 2001]. Following the procedure described therein, one obtains a tanglegram with 27 crossings and a taxon displacement of 42, matching the layout reported in the original study. Applying our DO-tanglegram algorithm to the same two trees, we obtain a tanglegram with only 2 crossings and a taxon displacement of 4 (see Fig. 2).

**Figure 2.**
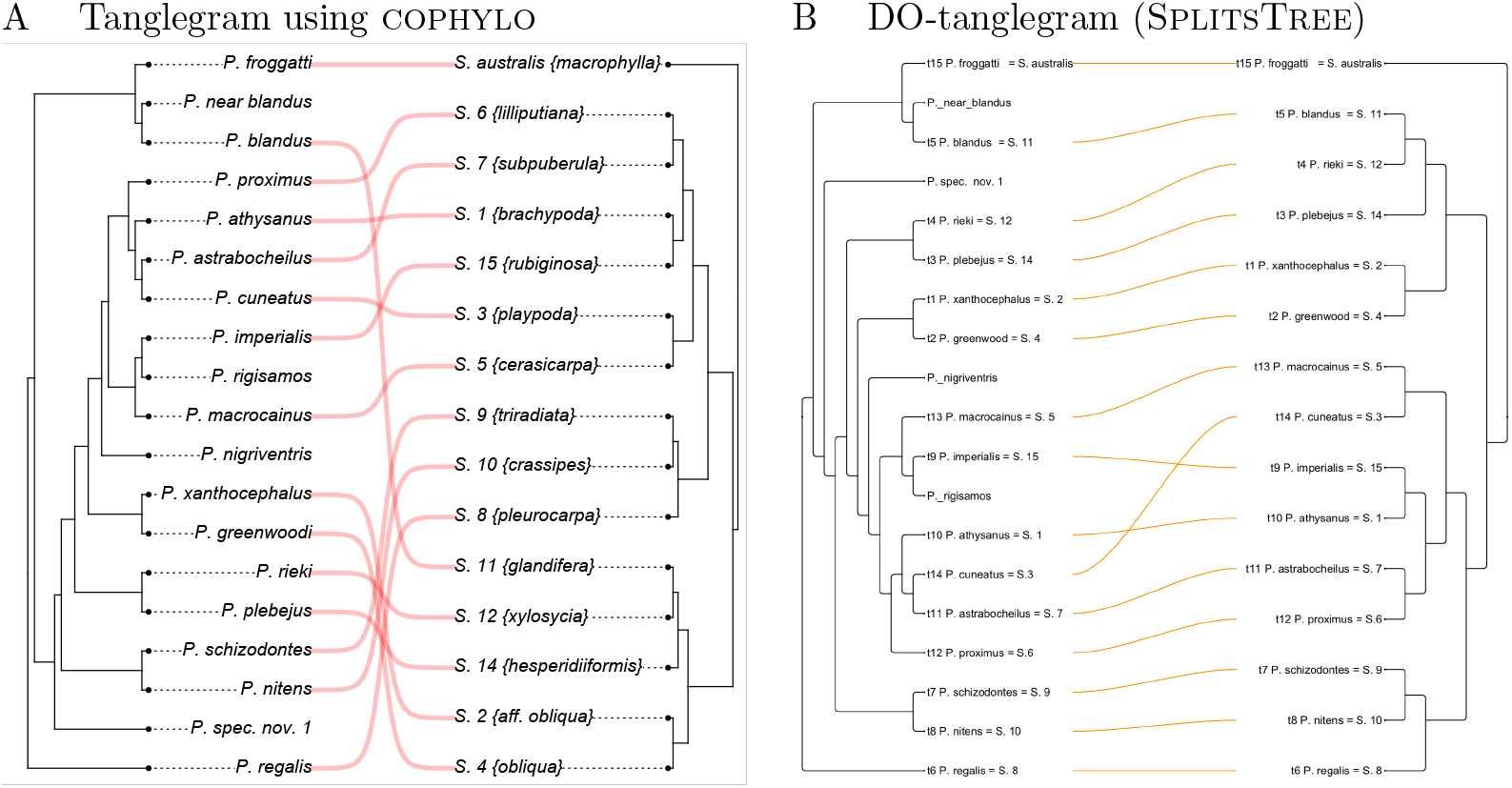
Comparison on wasp data. (A) Tanglegram obtained using the cophylo function in phytools, with 27 crossings and a taxon displacement of 42. This is the same layout as in [López-Vaamonde et al., 2001]. (B) Tanglegram obtained using the DO-tanglegram algorithm with NN presorting, with only 2 crossings and a taxon displacement of 4.

We also performed a systematic comparison on synthetic phylogenetic trees. To generate evaluation datasets, we began with a large background tree on 500 taxa. For a given set of parameters *n >* 0, *r* ≥ 0, *m* ∈ [0, 1], and *c* ∈ [0, 1], we first randomly extracted a subtree on *n* taxa from the background tree. Next, we created two input trees, *t*_1_ and *t*_2_, by independently applying *r* × *n* rooted subtree prune-and-regraft (rSPR) operations to the extracted tree. If *m >* 0, we randomly removed a proportion *m* of taxa from each tree. Similarly, if *c >* 0, we randomly contracted a proportion *c* of internal edges in each tree.

We constructed one pair of trees for each combination of *n* ∈ {50, 60, …, 450}, *r* ∈ {0.1, 0.2, 0.3}, *m* ∈ {0, 0.1}, and *c* ∈ {0, 0.1}. We ran both DO-tanglegram and cophylo (using the command `cophylo(t1, t2, rotate=TRUE)`) on every input pair and measured the total taxon displacement, number of crossings, and wall-clock runtime. On average, both methods required only a few seconds per dataset.

The results in Figure 3 indicate that our new method produces tanglegrams with lower taxon displacement on 99% of the datasets and with fewer inter-taxon crossings on 75% of the datasets, in comparison to cophylo.

**Figure 3.**
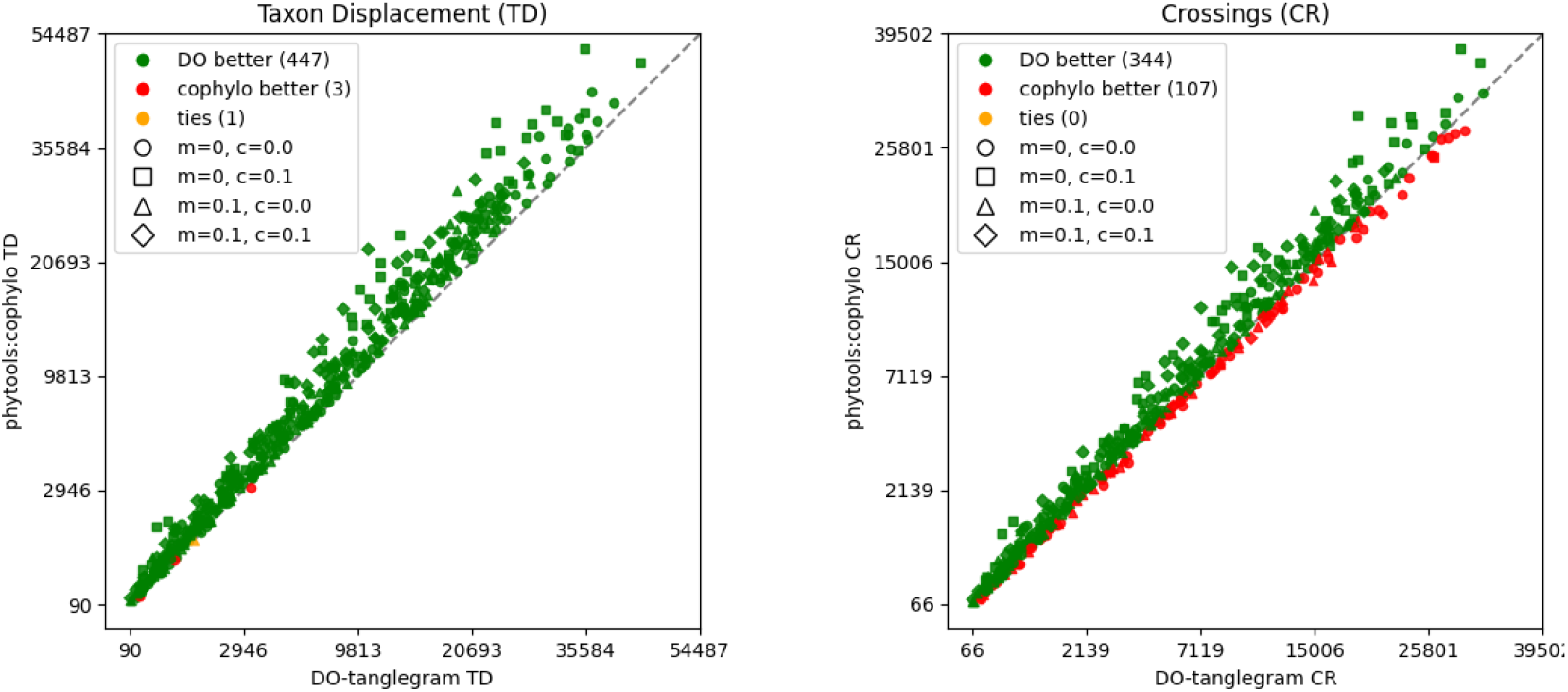
Comparison of DO-tanglegram and cophylo on pairs of synthetic phylogenetic trees, with 50 − 450 taxa, under varying proportions of reticulations, missing taxa (*m*), and contracted edges (*c*). On the left we compare taxon displacement and on the right we compare number of crossings.

### DO-tanglegrams on networks

The DO-tanglegram method is explicitly designed to work on rooted phylogenetic networks. Because it allows one to optimize both taxon displacement and reticulate displacement, and also supports the computation of one-sided tanglegrams, it represents an improvement over the current state-of-the-art method, NN-tanglegram [Scornavacca et al., 2011] as implemented in Dendroscope, which only allows two-sided computations and focuses only on optimizing the inter-taxon edge crossings.

In the paper introducing the NN-tanglegram method, the authors display a tanglegram between a rooted phylogenetic network and a rooted phylogenetic tree, based on data from [Kim and Donoghue, 2008]. In the NN-tanglegram layout, the taxon displacement is 128, number of crossings 90, and reticulate displacement 238. Using our DO-tanglegram heuristic on the same input, we obtain a tanglegram with taxon displacement 130, number of crossings 99, and reticulate displacement 93 (see Figure 4).

**Figure 4.**
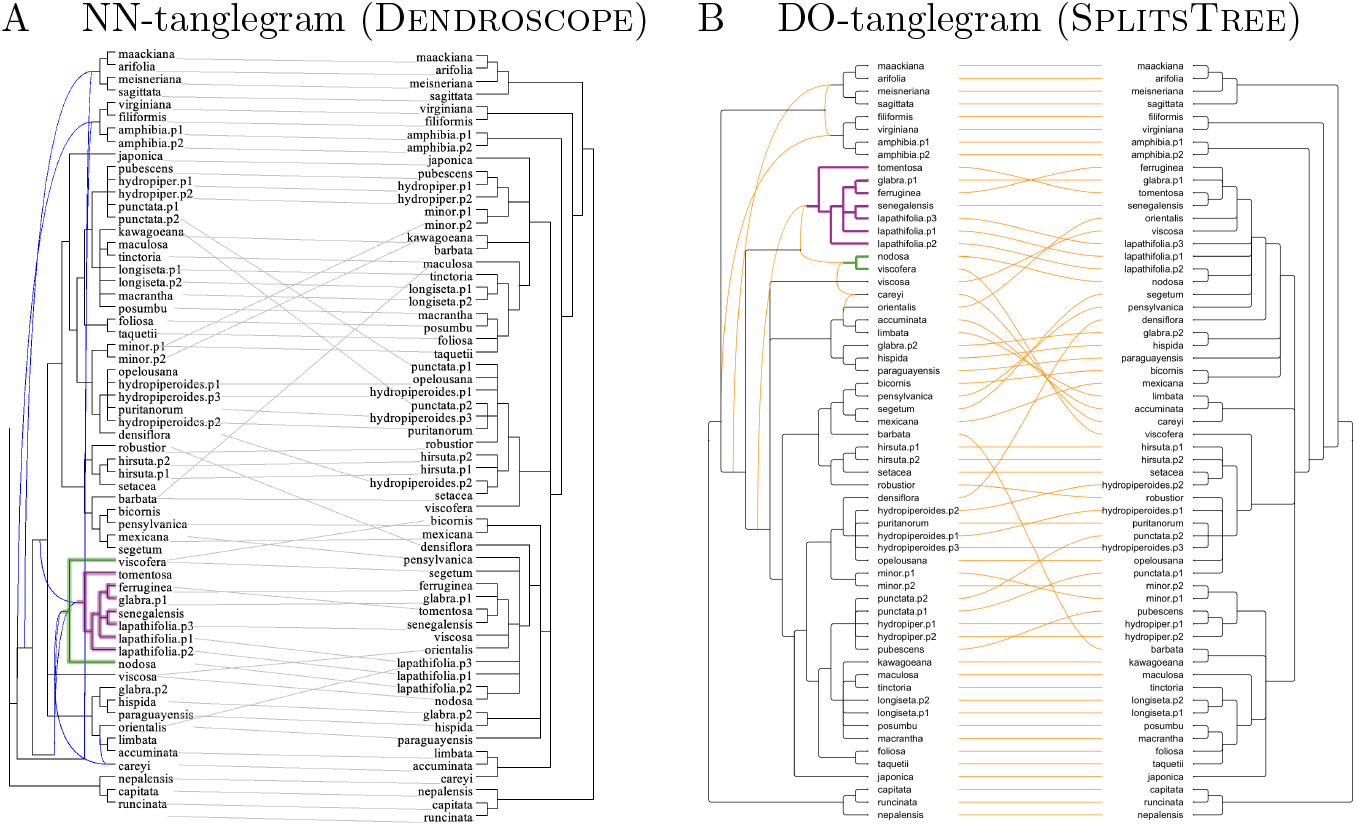
Comparison. (A) Tanglegram obtained using the NN-tanglegram method implemented in Dendroscope. (B) Tanglegram obtained using the DO-tanglegram. Note that (A) contains two nested trees (highlighted in green and purple), whereas the use of a backbone tree in the DO-tanglegram algorithm prevents this from happening in (B).

This example highlights an important conceptual difference between the two algorithms. In DO-tanglegrams, the leaves descending from a given parent or LSA-parent always appear as a contiguous block in the layout; they are never interleaved with leaves from other subtrees. Tanglegrams produced using the NN-tanglegam method do not necessarily satisfy this property. In the example, the two leaves of the green subtree are separated by leaves belonging to the purple subtree. While such a placement may reduce displacement measures, it does not necessarily improve the clarity of the visualization.

Here we use synthetic phylogenetic networks to compare the performance of the new DO-tanglegram algorithm, as implemented in SplitsTree, with the NN-tanglegram algorithm implemented in Dendroscope. To generate these evaluation datasets, we began with a large background tree on 500 taxa. For a range of target sizes (50–400, in steps of 10) and for each choice of *m* ∈ {0, 0.05} and *c* ∈ {0, 0.05}, we extracted two identical subtrees of the specified size from this background tree. In each subtree, we then randomly and independently removed a proportion *m* of all taxa and contracted proportion *c* of all internal edges, thus also considering datasets with missing taxa and unresolved nodes. Finally, in both trees, we randomly and independently added ten new edges to introduce reticulations.

Figure 5 compares the resulting taxon displacement and total reticulate displacement for the two algorithms. With respect to taxon displacement, DO-tanglegram often performs better than NN-tanglegram, especially on datasets with missing taxa. DO-tanglegram achieves lower total reticulate displacement in over 80% of all cases. (We note that the NN-tanglegram implementation crashed on ten input pairs; these were omitted from the analysis.) The mean wall-clock time required on these samples was 25 seconds (35 standard deviation) for DO-tanglegram and 102 seconds (262 seconds standard deviation) for NN-Tanglegram, suggesting a good speedup of the new method versus the old one.

**Figure 5.**
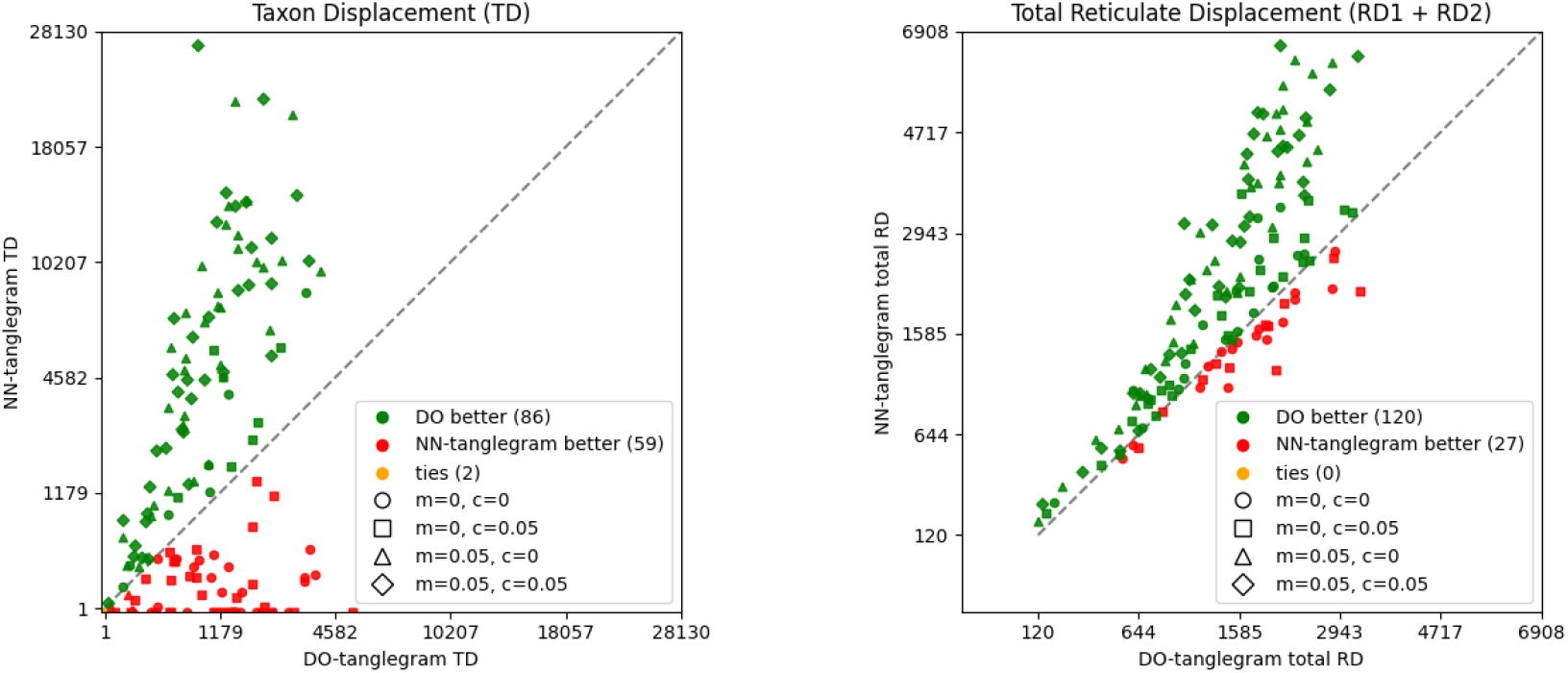
Comparison of DO-tanglegram (SplitsTree) and NN-tanglegram (Dendroscope) on synthetic pairs of rooted phylogenetic networks, on 50–400 taxa, each with 10 reticulations, under varying proportions of missing taxa (*m*) and contracted edges (*c*). Left: taxon displacement. Right: total reticulate displacement.

## 3 Discussion

Tanglegrams are used in biological research for comparing phylogenetic trees. Moreover, there is much research focused on developing methods for computing phylogenetic networks, whose aim is to explicitly represent reticulate evolutionary processes such as speciation-by-hybridization, horizontal gene transfer, and reassortment. Hence, there is a need for versatile approaches capable of computing tanglegrams between phylogenies that may be trees or rooted networks (with both combining and transfer reticulations), and that may contain unresolved nodes and missing taxa.

In this work we introduced the DO-tanglegram approach. The key idea is to jointly minimize both taxon displacement and reticulate displacement. Our results demonstrate that the method performs significantly better than state-of-the-art approaches—cophylo for trees and the NN-tanglegram algorithm in Dendroscope for networks. An implementation of the algorithm is provided in SplitsTree.

The NN-tanglegram method focuses exclusively on minimizing inter-taxon crossings and can only be applied in a two-sided setting. In contrast, the DO-tanglegram algorithm is more general: it explicitly optimizes both taxon displacement and reticulate displacement, and it supports both one-sided and two-sided tanglegram computations.

As datasets grow larger and more complex, with increased taxon sampling, missing data, and higher levels of reticulation, interpreting relationships among phylogenies becomes more challenging. By explicitly minimizing both taxon and reticulate displacement, DO-tanglegrams aim to produce layouts that preserve structural correspondence and reduce visual confusion. Such improvements can facilitate comparative studies in areas including gene tree–species tree discordance, host–parasite coevolution, and the analysis of hybridization and horizontal gene transfer, where understanding relationships between multiple phylogenetic hypotheses is essential.

Although the DO-tanglegram method performs well on the examples considered here, several limitations should be acknowledged. The quality of a resulting tanglegram may depend on the extent of taxon overlap and the degree of discordance between the phylogenies under comparison. In cases of extreme structural divergence, no tanglegram representation may adequately convey all relevant evolutionary signal, and alternative visualizations—such as agreement subtrees, splits graphs, or reconciliation models—may be more appropriate. Furthermore, while the heuristic is efficient for the dataset sizes tested here, its performance on substantially larger or highly reticulated networks remains to be systematically tested.

## 4 Materials and Methods

We adopt the same basic definitions as in [Huson, 2025]. Let *X* be a set of taxa. A rooted phylogenetic network *N* = (*V, E, ρ, λ*) on *X* is a directed graph with node set *V*, edge set *E* ⊆ *V* × *V*, root node *ρ* ∈ *V*, and taxon labeling *λ*, such that [Huson et al., 2010]:

1. *N* is a directed acyclic graph (DAG),
2. there is exactly one root node with in-degree 0, namely *ρ* (so in particular *N* is connected),
3. the labeling *λ* : *X* → *V* is a bijection between *X* and the set of leaves (nodes of out-degree 0),
4. there are no “through nodes,” i.e. nodes with both in-degree 1 and out-degree 1, and
5. all leaves have in-degree at most 1.

A node *v* is called a *tree node* if it has in-degree ≤ 1, and a *reticulate node* otherwise. An edge *e* = (*v, w*) is called a *tree edge* or a *reticulate edge* depending on whether its target *w* is a tree node or a reticulate node, respectively. To ensure practical relevance, we explicitly allow nodes to be *multifurcating*, with out-degree greater than two, and *multicombining*, with in-degree greater than two.

If there are no reticulate edges, then *N* is a rooted phylogenetic tree. In this case, *N* is planar and can be drawn in the plane without edge crossings. In contrast, if reticulate edges are present, *N* may be non-planar, and any drawing of *N* may necessarily involve crossings.

We view rooted phylogenetic networks as a natural generalization of rooted phylogenetic trees. When drawing such a network, the goal is that *tree edges never cross each other*, while *reticulate edges may cross tree edges and reticulate edges*. To enhance visual clarity, one aims to reduce the number and extent of crossings, and in particular to avoid steeply slanted reticulate edges. To this end, Huson [2025] introduced the concept of *reticulate displacement* and described an algorithm for obtaining network layouts that minimize this quantity.

In the present work, we extend these ideas from single-network layouts to the problem of computing tanglegrams for two rooted phylogenetic networks *N*_1_ and *N*_2_, allowing nodes of arbitrary in-degree or out-degree, and allowing the two networks to be defined on different but overlapping taxon sets *X*_1_ and *X*_2_.

## 5 The backbone tree and reticulate displacement

Let *N* be a phylogenetic network on taxon set *X*, with root *ρ*.

Recall that there are two distinct ways to interpret reticulate nodes in a rooted phylogenetic network [Huson et al., 2010, Huson, 2025]: in a *combining view*, a reticulate node represents a *combining event*, such as *hybridization-by-speciation*, where all incoming edges are treated equally and rendered in a similar fashion (see Fig. 1B of Huson, 2025). In contrast, in a *transfer view* a reticulate node represents a *transfer event*, such as *horizontal gene transfer*. In this case, one incoming edge—the *transfer-acceptor edge*—represents the main lineage and is drawn like a regular tree edge, while all other incoming edges represent transferred material and are drawn as reticulate edges (see Fig. 1D of Huson, 2025).

Let *v* be a reticulate node. The following operation converts *v* into a tree node. If *v* is associated with a transfer event, then one of its incoming edges *e* has been declared the transfer-acceptor edge, and we delete all other incoming edges. Otherwise, *v* is associated with a combining event. In this case, determine the *lowest stable ancestor LSA*(*v*), which is the last node on all paths from the root *ρ* to *v*. Then replace all incoming edges of *v* by a single incoming edge from *LSA*(*v*).

Application of this operation to all reticulate nodes yields a tree with root *ρ* that has the same nodes as *N*, including all leaves. We call *B* = *B*(*N*) the *backbone tree* of *N*. Note that the backbone tree is not necessarily a proper phylogenetic tree, because it may contain unlabeled leaves and/or through nodes.

A basic left-to-right layout of the backbone tree *B* can be obtained by first assigning *x*-coordinate *x*(*v*) = *d*, where *d* is the number of edges on the path from the root *ρ* to node *v*. Then we set the *y*-coordinate *y*(*ℓ*) = *k* when *ℓ* is the *k*-th leaf encountered in a post-order traversal of the tree, and for an internal node *v* we set *y*(*v*) = avg_*ℓ<v*_{*y*(*ℓ*)}, the average *y*-coordinate of all leaves *ℓ* below *v*. See [Huson, 2025] for further elaboration.

The *reticulate displacement* of a left-to-right layout is defined as

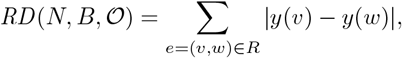

where *R* ⊆ *E* is the set of reticulate edges (i.e., combining or transfer edges). This measure depends solely on the *y*-coordinates assigned to nodes during the post-order traversal of the backbone tree *B*, which in turn depend on the ordering *O* in which the children of each node *v* are visited. To reduce visual clutter and avoid steeply slanted reticulate edges in a drawing of *N*, one should use an ordering *O* of *B* that minimizes the reticulate displacement [Huson, 2025].

## 6 One-sided tanglegram layout

In the one-sided tanglegram problem, the layout of one rooted phylogeny (tree or network) is fixed, and we seek a suitable layout for the other. Let *N*_1_ and *N*_2_ be two rooted phylogenies (either can be a tree or network) on taxon sets *X*_1_ and *X*_2_, respectively, and let *B*_1_ and *B*_2_ be their corresponding backbone trees. Assume that the layout (ordering) *O*_1_ of *B*_1_, and thus also the coordinates *x* and *y* of the nodes in *N*_1_, are fixed. Our goal is to determine a good ordering *O*_2_ for *B*_2_, yielding coordinates *x* and *y* for *N*_2_.

A good ordering will attempt to minimize the *reticulate displacement* of *N*_2_ and also the *taxon displacement* of the tanglegram, which we define as

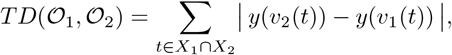

where *v*_1_(*t*) and *v*_2_(*t*) are the leaves of *N*_1_ and *N*_2_, respectively, that are labeled by taxon *t*.

Putting both together, we define the *one-sided tanglegram score* of a left-to-right layout of *N*_2_ relative to *N*_1_ as

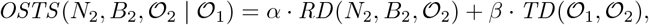

where *α, β* ∈ [0, 1] can be used to balance the avoidance of steep reticulation edges (first term) against the vertical misalignment of corresponding taxa (second term). A good one-sided layout for *N*_2_ is then given by any ordering *O*_2_ that minimizes this quantity.

In [Huson, 2025], we show that optimizing reticulate displacement is computationally hard, and introduce a heuristic based on a pre-order traversal of *B*_2_: for each node *v* that is the lowest stable ancestor (LSA) of some reticulate node, or is the source of a transfer edge, consider permutations of its children in *B*_2_ so as to minimize the total reticulate displacement. If the number of children is at most eight, all permutations are evaluated exhaustively; otherwise, a heuristic search is performed by iteratively swapping pairs of children, using simulated annealing [Kirkpatrick et al., 1983] to escape local minima. In contrast, if *α* = 0, then the objective reduces to minimizing only the taxon displacement for the backbone tree (reticulation edges are ignored), which can be solved in polynomial time [Venkatachalam et al., 2010].

To address the combined optimization of reticulate and taxon displacement, we propose to modify this heuristic search in two ways. First, at the above-mentioned nodes, we use *OSTS* rather than *RD* as the objective. Second, we also process all other interior nodes in a similar fashion, using the taxon displacement *TD* as the local objective function.

We refer to this as the *one-sided DO-tanglegram heuristic*.

This heuristic has an obvious weakness. For example, if *N*_2_ is a rooted star tree (a tree consisting only of a root and a set of leaves), then the ordering of the leaves is fully flexible and there exists an optimal layout with zero taxon displacement. However, if the out-degree of *ρ* exceeds the heuristic threshold beyond which only a subset of orderings is considered, then the optimal layout may be missed.

To mitigate this, we define the *rank* of *t* ∈ *X*_1_ as the *y*-coordinate of the associated leaf in *N*_1_. Using a post-order traversal, for each node *u* of the backbone tree *B*_2_, we determine the taxon *t* ∈ *X*_1_ ∩ *X*_2_ of smallest rank that labels a leaf descendant of *u*. Now, when considering permutations of the children of some node *v* in the heuristic, we will always first consider the permutation obtained by ordering the children of *v* according to the smallest-rank taxa associated with their subtrees, leaving children without associated taxa fixed.

### Accommodating missing taxa

Let *X* = *X*_1_ ∩ *X*_2_ denote the set of taxa shared by the two networks (or trees), and let *n* = |*X*|. Since *N*_1_ and *N*_2_ may each contain additional taxa not present in the other, we compute taxon displacement based only on *X*, using ranks rather than raw *y*-coordinates.

In the one-sided problem, the ordering *O*_1_ of the backbone tree *B*_1_ is fixed. This determines a ranking *π*_1_ : *X* → {1, …, *n*} of the shared taxa, obtained by traversing *B*_1_ in post-order and recording the order in which the shared leaves are visited. For any candidate ordering *O*_2_ of the backbone tree *B*_2_, we obtain a corresponding ranking *π*_2_ : *X* → {1, …, *n*} in the same way.

We then define the *taxon displacement* between *O*_1_ and *O*_2_ as

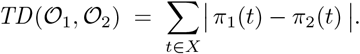

## 7 Two-sided tanglegram layout

In the two-sided tanglegram layout problem, the layout of neither network is fixed, and both can be modified in order to obtain a good overall drawing. Let *N*_1_ and *N*_2_ be two rooted networks (or trees) on taxon sets *X*_1_ and *X*_2_, respectively, and let *B*_1_ and *B*_2_ be their corresponding backbone trees.

The goal is to determine good orderings *O*_1_ for *B*_1_ and *O*_2_ for *B*_2_, yielding coordinates *x* and *y* for all nodes in *N*_1_ and *N*_2_.

We evaluate layouts using the *two-sided tanglegram score*, defined as

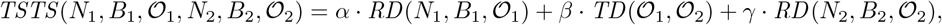

where the coefficients *α, β, γ* ∈ [0, 1] balance the competing goals of reducing reticulate displacement in each network (first and last terms) and reducing taxon displacement between the two sides (middle term).

Minimizing *RD* on either network is computationally hard [Huson, 2025]. While minimizing *TD* alone can be done in polynomial time, minimizing the number of inter-taxon edge crossings in the two-sided case is NP-complete, even when both *N*_1_ and *N*_2_ are binary trees [Fernau et al., 2010].

To address this problem heuristically, we apply the one-sided DO-tanglegram heuristic alternately to *N*_1_ and *N*_2_: in each step we optimize the layout of one network while keeping the other fixed, then switch roles. Iterating this process yields a pair of orderings (*O*_1_, *O*_2_) that together define a joint layout [Bansal et al., 2009]. We refer to this as our *two-sided DO-tanglegram heuristic*.

### Presorting using neighbor-net

In a presorting step, the NN-tanglegram method implemented in Den-droscope uses the neighbor-net algorithm [Bryant and Moulton, 2004] to compute an initial circular ordering of the taxa present in both networks. It then applies a greedy heuristic that repeatedly swaps subnetworks in order to reduce the number of inter-taxon crossings. We incorporate the same presorting strategy in our approach and refer to it as the *NN-presorting heuristic*.

Recall that every tree node *v* of a rooted phylogenetic network induces a non-empty cluster, namely the set of taxa labeling the leaf descendants of *v*, the so-called *hardwired clusters* [Huson et al., 2010]. Let *C* be the set of all hardwired clusters extracted from *N*_1_ and *N*_2_, restricted to the set *X* of taxa common to both networks, and of size at least two. For every pair of taxa *i, j* ∈ *X* we define their distance as

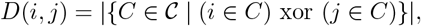

that is, the number of clusters in *C* that contain exactly one of *i* and *j*.

We run the agglomerative phase of the neighbor-net algorithm on the distance matrix *D* to obtain a circular ordering *O* of *X*, and then use this ordering to improve the initial layout of the backbone trees *B*_1_ and *B*_2_. Let the *rank* of a taxon *t* ∈ *X* be its position in *O*. For each of *B*_1_ and *B*_2_, we perform a post-order traversal and, at every internal node *v*, sort its children by the increasing average rank of the taxa in their subtrees.

This presorting step aims to exploit the cluster structure of the two phylogenies in order to derive a taxon order that is as consistent as possible between the networks. If both inputs are trees and their cluster sets are fully compatible, this preprocessing guarantees a solution with zero inter-taxon edge crossings [Scornavacca et al., 2011]. In general, however, NN-presorting simply provides an informed initialization for the DO-tanglegram heuristic: in many cases it substantially reduces crossings and displacement, while in others it may lead to less favorable layouts due to the heuristic nature of the subsequent optimization.

### Advanced settings

The heuristic search is executed in parallel using 32 jobs (the default), each initialized with a different random ordering of the children below every LSA node, in the case of a network, or below all children, in the case of a tree. This degree of parallelization adds little to the overall wall-clock time and gives rise to improved layouts.

In our heuristic search, the default parameters for simulated annealing are a start temperature of 1000, an end temperature of 0.01, 1000 iterations per temperature step, and a cooling rate of 0.95. Although these settings are not exposed directly in the SplitsTree user interface, they—together with the default number of parallel jobs for network computations—can be adjusted through the application preferences, as described in the online SplitsTree user manual.

## Availability

The algorithm is implemented in our open source program SplitsTree, available here: https://github.com/husonlab/splitstree6.

All data used to produce the figures are available here: https://github.com/husonlab/splitstree6/tree/main/tanglegram-paper.

